# Phenotypic, genomic, and transcriptomic heterogeneity in a pancreatic cancer cell line

**DOI:** 10.1101/2022.11.11.516211

**Authors:** Gengqiang Xie, Liting Zhang, Olalekan H Usman, Sampath Kumar, Chaity Modak, Dhenu Patel, Megan Kavanaugh, Xian Mallory, Yue Julia Wang, Jerome Irianto

## Abstract

**Objectives:** To evaluate the suitability of the MIA PaCa-2 cell line for studying pancreatic cancer intratumor heterogeneity, we aim to further characterize the nature of MIA PaCa-2 cells’ phenotypic, genomic, and transcriptomic heterogeneity.

**Methods:** MIA PaCa-2 single-cell clones were established through flow cytometry. For the phenotypic study, we quantified the cellular morphology, proliferation rate, migration potential, and drug sensitivity of the clones. The chromosome copy number and transcriptomic profiles were quantified using SNPa and RNA-seq, respectively.

**Results:** Four MIA PaCa-2 clones showed distinctive phenotypes, with differences in cellular morphology, proliferation rate, migration potential, and drug sensitivity. We also observed a degree of genomic variations between these clones in form of chromosome copy number alterations and single nucleotide variations, suggesting the genomic heterogeneity of the population, and the intrinsic genomic instability of MIA PaCa-2 cells. Lastly, transcriptomic analysis of the clones also revealed gene expression profile differences between the clones, including the uniquely regulated *ITGAV*, which dictates the morphology of MIA PaCa-2 clones.

**Conclusions:** MIA PaCa-2 is comprised of cells with distinctive phenotypes, heterogeneous genomes, and differential transcriptomic profiles, suggesting its suitability as a model to study the underlying mechanisms behind pancreatic cancer heterogeneity.

## Introduction

In most cancers, the transformed cells undergo cycles of genetic diversification, selection of adaptive clones, and clonal expansion, a process that mirrors Darwin’s natural selection ^1^. Specific mutations provide a growth advantage to a subpopulation of cells, giving rise to the evolution of distinctive clones ^2^. The evolution process will then lead to a tumor comprised of cells with heterogeneous phenotypes, including response to therapy ^3^. This is certainly the case for pancreatic ductal adenocarcinoma (PDAC), the most common type of pancreatic cancer. Intratumor genomic heterogeneity within PDAC tumors has been reported by several studies ^4–7^, suggesting the genomic instability of PDAC cells. Moreover, two subtypes of PDAC cells were revealed by multiple transcriptomic studies ^8–11^: the more common classical/progenitor subtype and the more aggressive basal-like/squamous subtype. Interestingly, several recent studies have shown the existence of these subtypes within a PDAC tumor ^12–15^, suggesting the plasticity of PDAC cells and the phenotypic heterogeneity within the tumor, which might be driven by the genomic events during tumor evolution ^12,13^. However, the underlying mechanisms behind PDAC genomic instability and the observed phenotypic and genomic heterogeneity are largely unknown. One possible approach to uncovering the underlying mechanisms of PDAC heterogeneity is to perform a mechanistic study using PDAC cell lines that show phenotypic and genomic heterogeneity.

MIA PaCa-2 is one of the commonly used primary tumor-derived PDAC cell lines ^16^. Genomically, MIA PaCa-2 has mutations within the three most frequently mutated genes in PDAC ^17^: G12C in *KRAS*, R248W in *TP53*, and homozygous deletion of *CDKN2A*. In addition, transcriptionally, MIA PaCa-2 cells have been categorized as the basal-like/squamous subtype ^8^. By isolating single-cell clones, several studies have shown that MIA PaCa-2 comprises cells with differential phenotypes in multiple aspects, including adhesion, integrin expression, anchorage-independent growth, and invasion ^18,19^, suggesting the phenotypic heterogeneity of MIA PaCa-2 cells. Based on these findings, MIA PaCa-2 may be a good candidate on which to perform a mechanistic study of PDAC heterogeneity. In this study, to evaluate its suitability for a mechanistic study, we aimed to further characterize the nature of MIA PaCa-2 phenotypic, genomic, and transcriptomic heterogeneity. First, we established MIA PaCa-2 clones through single-cell flow cytometry and chose four clones that display clear phenotypic differences in their cellular morphology, proliferation rate, and migration potential. Then, using a single-nucleotide polymorphism microarray (SNPa), we observed genomic heterogeneity between the MIA PaCa-2 clones in the form of chromosome copy number alterations. Our RNA-seq analysis also showed significant differences between the transcriptomic profiles of the clones. Lastly, from the transcriptomic analysis, we identified a gene, *Integrin Subunit Alpha V* (*ITGAV*), that is uniquely regulated between the clones and its modulation dictates MIA PaCa-2 cell morphology, suggesting a possible driver of the observed differences in cellular morphology, linking the phenotypic and transcriptional heterogeneity.

## Materials and Methods

### Cell culture

The MIA PaCa-2 human pancreatic cell line was obtained from ATCC. Cells were maintained in Dulbecco’s modified Eagles medium (DMEM) supplemented with 10% fetal bovine serum (Genesee Scientific) and 1% penicillin-streptomycin (Sigma-Aldrich). MIA PaCa-2 clones were generated by single-cell flow cytometry sorting at Florida State University’s Flow Cytometry Laboratory. Briefly, MIA PaCa-2 cells were harvested after dissociation with trypsin (Corning) at 37°C for 5 minutes. Cells were washed once with PBS and resuspended with 1 ml of PBS supplemented with 10% fetal bovine serum and 1% penicillin-streptomycin. Cell suspensions were filtered through a BD FACS tube with a 70 μm strainer top. Single cells were sorted using the FACSCanto instrument (BD) by forward scatter to a 96-well plate (Genesee Scientific). Twenty-one MIA PaCa-2 single-cell clones were established after approximately 3 weeks of culture, and four of the 21 clones were expanded and used in this study. The same method was used to derive the secondary single-cell clones from the four primary single-cell clones. *ITGAV* and *ITGB5* overexpression was achieved by 24-hour transfection with Lipofectamine 2000 (ThermoFisher); the mApple-Alpha-V-Integrin-25 was a gift from Michael Davidson (Addgene plasmid # 54866; http://n2t.net/addgene:54866; RRID: Addgene_54866), the pCX-EGFP beta5 integrin receptor ^20^ was a gift from Raymond Birge (Addgene plasmid # 14996; http://n2t.net/addgene:14996; RRID:Addgene_14996). For *ITGAV* depletion, three gRNAs were selected by using CRISPick ^21,22^ and cloned into lentiCRISPR v2 plasmid ^23^, a gift from Feng Zhang (Addgene plasmid # 52961; http://n2t.net/addgene:52961; RRID: Addgene_52961). Lentiviral particles were produced through the transfection of the transfer plasmid, lentiCRISPR v2, and the packaging plasmids, psPAX2 (a gift from Didier Trono, Addgene plasmid # 12260; http://n2t.net/addgene:12260; RRID: Addgene_12260) and pCMV-VSV-G ^24^ (a gift from Bob Weinberg, Addgene plasmid # 8454; http://n2t.net/addgene:8454; RRID: Addgene_8454), into the HEK293T cells by using the TransIT-Lenti transfection reagent (Mirus Bio). Lentiviral transduction of the CRISPR construct was performed with 10 µg/mL polybrene (ThermoFisher).

### Microscopy and image processing

Images were taken either with an Olympus IX71 with an sCMOS camera (Prime 95B, Photometrics) or EVOS microscope (ThermoFisher), with 10x/0.3NA, 20x/0.75NA, or 40x/0.6NA objective. All the image analysis was done with ImageJ ^25^. Circularity and aspect ratio were measured using the built-in algorithm in ImageJ. For the circularity, the area and perimeter of the region of interest are measured, and the circularity is calculated based on the following formula: 4π*area/perimeter^2^. A value of 1 indicates a perfect circle, while as the value approaches 0, it indicates an increasingly elongated shape. For the aspect ratio, an ellipse is fitted to the region of interest, and the aspect ratio is calculated as the ratio between the major and minor axis of the ellipse. The increasing aspect ratio value indicates an increasingly elongated shape.

### Scratch wound healing assay

Cell migration was evaluated by a scratch wound healing assay as previously described ^26,27^. MIA PaCa-2 clone cells were seeded at a concentration of 100,000 cells/cm^2^ (80% confluent) in 12-well plates and cultured under normal culture medium until reaching fully confluent. 2 mM thymidine was introduced to the cells to block cell proliferation for 24 hours. Next, each well was artificially wounded in the middle by creating a scratch with a 1000 µL tip plastic pipette on the cell monolayer. Each well was washed once with PBS to remove the floating cells and then cultured under a normal culture medium supplemented with 2 mM thymidine until the end of the assay. Images of scratch wounds were captured using an EVOS microscope at 0, 24, and 48 hours after the scratch. The wound closure areas were measured with ImageJ ^25^ after 24 and 48 hours. To avoid scratch area variation, the relative wound closure area was calculated as [%] = (wound closure area^T0^ - wound closure area^T48^) * 100 [%]/wound closure area^T0^.

### Gemcitabine treatment

Cells were plated in a regular white-bottom 96-well plate and cultured for at least 24 hours prior to the gemcitabine treatment (MilliporeSigma). After 72 hours of gemcitabine treatment, the cell number was measured using the luminescence cell viability assay, as described above. To derive the gemcitabine dose-response curve, the luminescence data was fitted with the following equation ^28^:

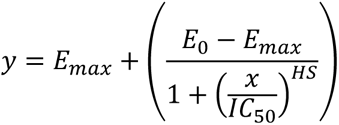

where *x* is the gemcitabine concentration, *y* is the corresponding luminescence readout, *E_0_* and *E_max_* are the top and bottom asymptotes of the curve, respectively, *IC_50_* is the gemcitabine concentration at the half-maximal effect, and *HS* is the Hill slope coefficient. The Hill slope coefficient was constrained to −1 to assume that gemcitabine binding follows the law of mass action. Curve fitting and calculation of IC50 and E_max_ were done using GraphPad Prism v.10 (GraphPad).

### Western blot

Western blotting was performed using standard methods. Briefly, cell pellets were harvested and resuspended in RIPA buffer with the addition of a phosphatase inhibitor cocktail as 50 µL buffer per million cells. Cell suspensions were then sonicated for 15 seconds and immediately put on ice for 30 minutes and cleared by centrifugation for 30 minutes at maximum speed. 10–15 µL of protein lysis were run per lane of 4–12% Bis-Tris gels (Thermo Fisher Scientific) using MOPS buffer. Proteins were transferred to nitrocellulose membrane at 20 V for 7 minutes using the iBlot™ 2 Transfer Stack (Thermo Fisher Scientific). Membranes were blocked in 5% non-fat milk powder in TBS+0.1% Tween (TBST) and incubated overnight at 4°C in primary antibody diluted in TBS solution at 1 µg/mL. Primary antibodies used were rabbit monoclonal antibody against ITGAV (ab179475, Abcam) and mouse monoclonal antibody against β-Actin (sc-47778, Santa Cruz Biotechnology). Membranes were incubated with corresponding horseradish peroxidase-conjugated second antibodies at room temperature for 1 hour. After three washes in TBST, membranes were exposed to ChromoSensor^TM^ One-Solution (GenScript) for 3–5 minutes. Membranes were quickly washed three times with water and imaged.

### Single nucleotide polymorphism microarray

DNA isolation was performed using the Blood and Cell Culture DNA Mini Kit (QIAGEN). The isolated DNA samples were sent to the Center for Applied Genomics at the Children’s Hospital of Philadelphia for the SNP array analysis using the Infinium Global Screening Array-24 v3.0 Kit (Illumina), which enabled the sequencing of 759,993 SNPs across the genome, at the average gap of around 4kb between SNPs. The chromosome copy number analysis was performed in GenomeStudio v.2.0 (Illumina) by using the cnvPartition v.3.2.1 plugin. The chromosome copy number calls were averaged every 1 Mb to derive the copy number profiles at 1 Mb resolution. Heatmaps were plotted in R by using the algorithm gtrellis v.1.28.0 ^29^. The hierarchical clustering was done using the heatmap.2 function from ggplots v.3.1.3 ^30^ in R.

For the SNP-based phylogenetic tree analysis, we transformed the SNP genotype into a trinary representation, classifying each SNP as alternative, non-alternative, or missing. Subsequently, we calculated pairwise distances between samples based on their respective genotypes. The distance between a pair of samples is calculated by the percentage of mismatches SNPs genotypes, out of the total 759,993 SNPs sequenced in the array. In light of a notable missing rate, we chose to exclude SNP sites with missing values in either sample of the pair. It is crucial to underscore that the excluded SNP sites differ for each pair of samples. Following the computation of pairwise distances, we applied the Neighbor-Joining algorithm for phylogenetic tree construction. This algorithm, a well-established method in evolutionary biology, iteratively merges pairs of nodes with the smallest pairwise distances until a comprehensive tree is formed. To address any potential issues with negative branch lengths, we implemented a correction by setting negative branch lengths to zero and transferring the difference to the adjacent branch lengths. This ensures that the total distance between adjacent pairs of terminal nodes remains unaltered and does not affect the overall topology of the tree ^31^.

### RNA-seq sample preparation and analysis

RNA isolation from the MIA PaCa-2 clones was performed by using the RNeasy Plus Micro Kit (QIAGEN). Libraries for RNA-seq were made with the NEBNext Ultra II Directional RNA Library Prep Kit for Illumina (NEB) per the manufacturer’s instructions, followed by 150-bp paired-end sequencing with the Novaseq 6000 at Florida State University’s Translational Science Laboratory, resulting in ∼25,000,000 reads for each sample. The quality of the fastq reads from the sequencing was examined with FastQC (version 0.11.9) ^32^, and aligned to the hg38 human reference genome using STAR v.2.7.9a ^33^. The aligned reads were scaled and normalized with the median of ratios using the DESeq2 v.1.32.0 ^34^. Differential expression was calculated using the result function in DESeq2, with the threshold of adjusted *p*-value < 0.05 and abs(log_2_FC) > 0.58. To derive the uniquely regulated genes of a given primary clone, three pair-wise differential expression analyses were done between the clone of interest and the other three clones. A gene is considered to be uniquely regulated when it is regulated in the same direction within all of the three analyses. For example, Clone #1 RNA-seq data was compared to the data of Clone #2, #3, and #4. A gene is considered uniquely upregulated only if the gene is upregulated in all comparisons: #1 vs #2, #1 vs #3, and #1 vs #4. The heatmap visualization and hierarchical clustering were done using the heatmap.2 function from ggplots v.3.1.3 ^30^ in R. The gene set enrichment analysis of the regulated genes was done using Enrichr ^35^, against the “hallmark” gene sets of the human molecular signatures database (MSigDB)^36^. The significant enrichment was selected through a threshold of adjusted *p*-value < 0.05 (FDR<0.05).

## Results

### Morphological heterogeneity of MIA PaCa-2 clones

To study whether heterogeneity exists in MIA PaCa-2 cells, we first flow-sorted a single cell into each well of a 96-well plate by forward scatter flow cytometry and expanded the cells to establish single-cell clones. Twenty-one single-cell clones were successfully established through this method. Interestingly, four distinct cellular morphologies were observed among the clones (**Figure 1A** and **S1**), named clones #1, #2, #3, and #4. Both clone #1 and clone #2 cells had a spindle-like morphology; however, clone #1 cells appeared to have a lower degree of cell spreading, resulting in a more elongated morphology. Clone #3 cells were cuboidal, whereas clone #4 cells were rounded and looked much smaller than cells from the other three clones. Almost half of the established clones have clone #1-like morphology, while the rest of the population consists of the other cell morphology types at more or less equal proportions (**Figure 1B**). We then quantified the cellular morphology of these four clones through three cell morphometric parameters (cell area, circularity, and aspect ratio in **Figure 1C**). Indeed, the cell surface area and the aspect ratio of clone #4 were significantly lower than the other three clones, whereas its circularity was significantly higher, reinforcing our observation of the rounded morphology of the clone #4 population. Likewise, the aspect ratio of clone #1 was higher than clone #2, confirming the more elongated morphology of clone #1 cells. These results demonstrate morphological heterogeneity within MIA PaCa-2 cells. Considering that cellular morphology is a gross phenotype that depends on many biological processes, such as integrin-ECM interaction^37^, focal adhesion maturation ^38^, cytoskeletal organization ^39^, and cellular polarity regulation ^40,41^, we anticipate that the variations in cellular morphology observed in this study serve as a reliable marker for heterogeneity within the parental bulk population. Consequently, we selected the four clones to conduct a more in-depth investigation of population heterogeneity across phenotypic, genomic, and transcriptomic levels.

**Figure 1.**
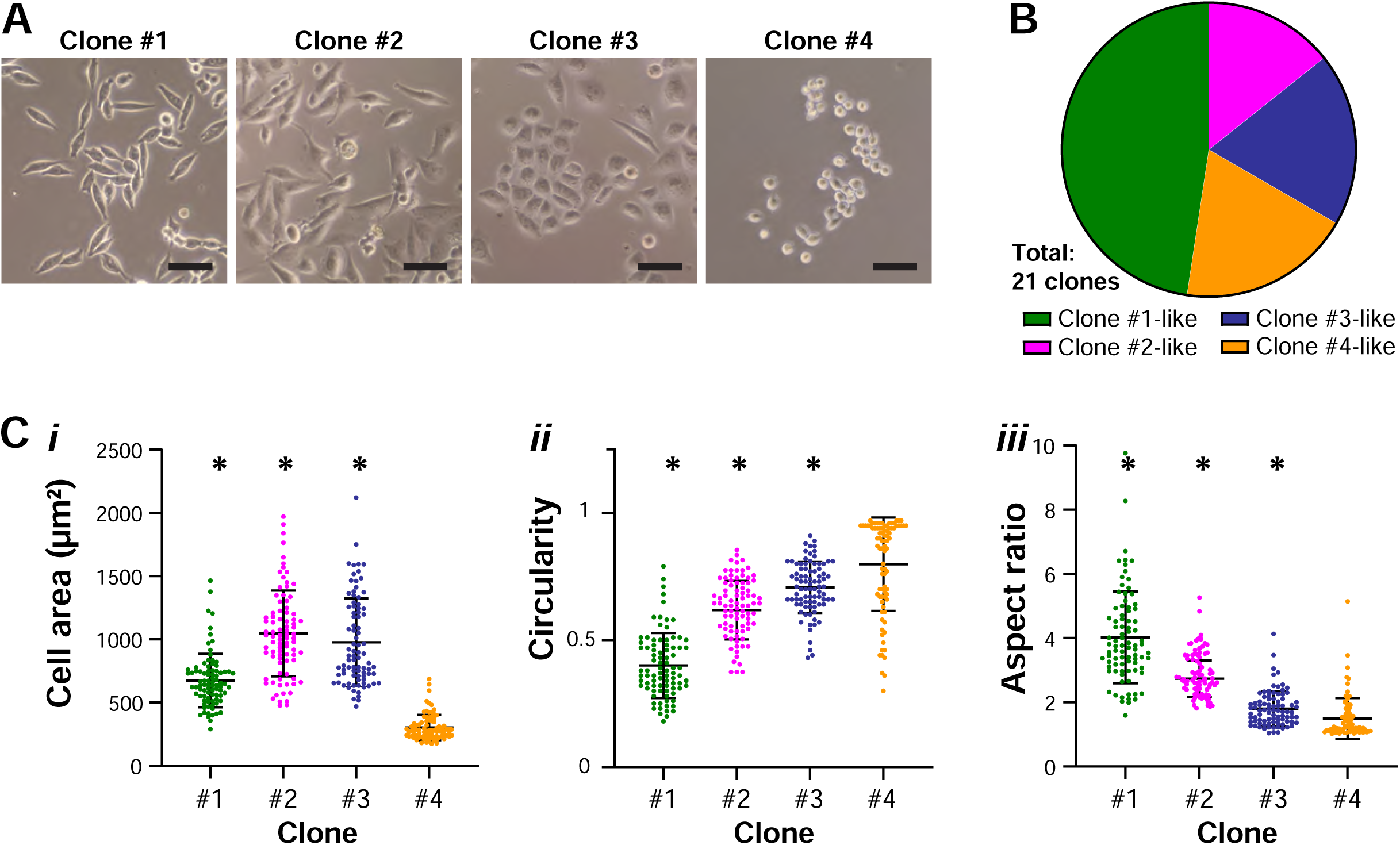
Morphological heterogeneity of MIA PaCa-2 clones. (**A**) Pie chart showing the proportion of MIA PaCa-2 single-cell clones based on their cellular morphology. (**B**) Representative images of the four clones with distinctive cellular morphology (scale bar = 50 μm). (**C**) Three cell morphometric parameters (cell area, circularity, and aspect ratio) were analyzed by ImageJ, showing significant differences against the rounded clone #4 (N = 3 experiments, Welch’s t-test, *p* < 0.05 vs. clone #4).

### Proliferation rate, migration potential, and gemcitabine sensitivity of MIA PaCa-2 clones

We then investigated whether the MIA PaCa-2 clones show other phenotypic differences, such as proliferation rate, migration potential, and drug sensitivity. First, we seeded the same number of cells from the four MIA PaCa-2 clones at day 0, cultured them for 4 days, and imaged them every day from day 1 to day 4. Cell numbers were counted each day for each of the clones. We found that clone #1 was the fastest-proliferating cell line, while clone #3 was the slowest (**Figure 2A–B**). When compared with clone #4, the other three clones displayed significant differences on day 4. The result suggests a differential proliferation rate between the MIA PaCa-2 clones.

**Figure 2.**
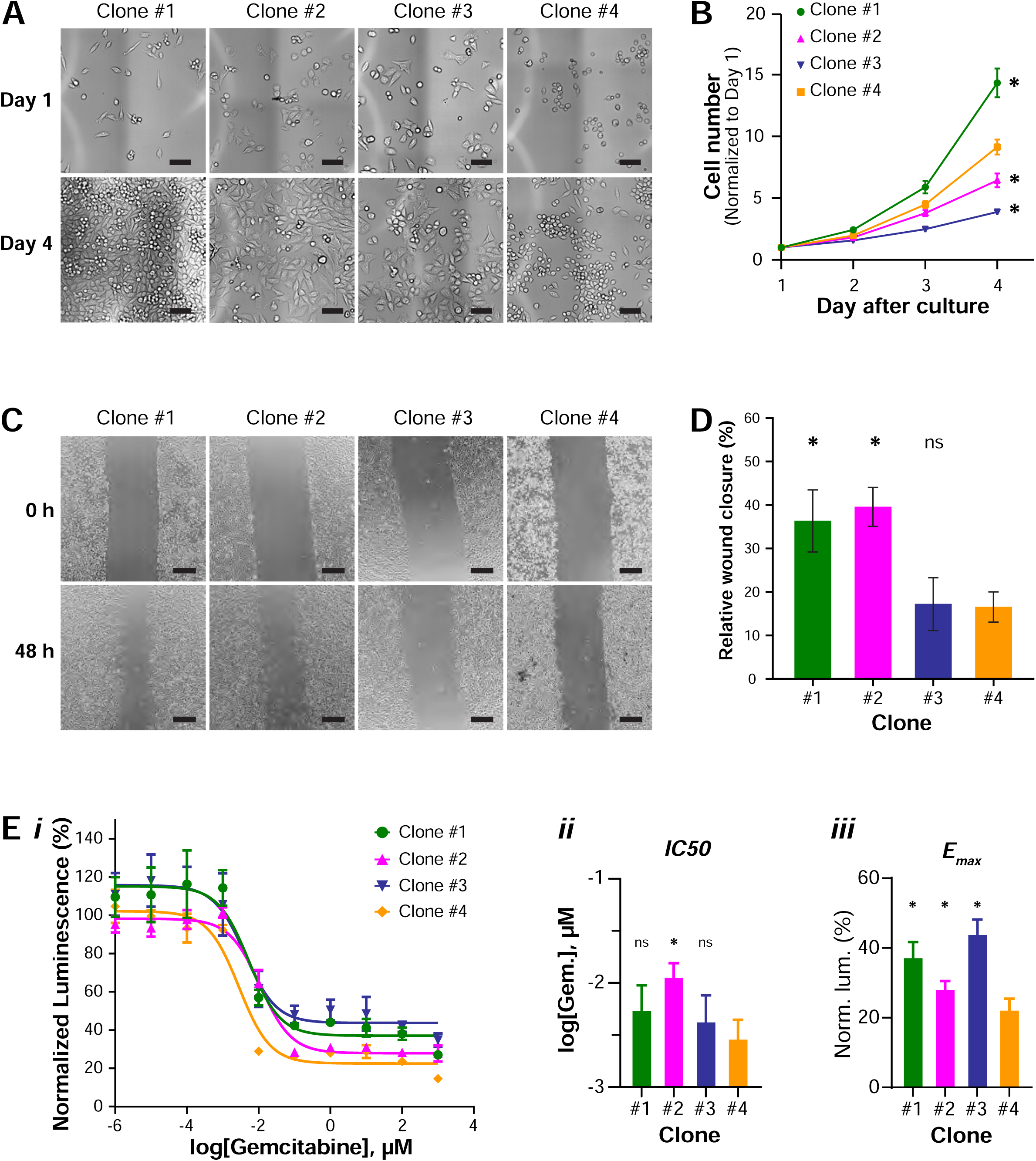
Differential proliferation, migration rate, and drug sensitivity of MIA PaCa-2 clones. (**A**) Representative images of the MIA PaCa-2 clone cells on day 1 and day 4 after seeding (scale bar = 100 μm). (**B**) The plot of cell numbers of different MIA PaCa-2 clones over a 4-day culture period. The cell numbers were normalized to the cell number of day 1 in each of the clones (N = 3 experiments, Welch’s t-test, *p* < 0.05 vs. clone #4). (**C**) Representative images of the scratch wound healing assay of the four MIA PaCa-2 clones at zero and 48 hours after the scratch (scale bar = 300 μm). (**D**) The plot of the relative wound closure of the different MIA PaCa-2 clones at 48 hours (N = 3 experiments, Welch’s t-test, *p* < 0.05 vs. clone #4). (**E**) MIA PaCa-2 clones #1, #2, #3, and #4 were treated with gemcitabine for 72 hours and the cell numbers were measured using a luminescence cell viability assay. (***i***) Curve fitting was performed on the luminescence data to derive the dose-response curves and the corresponding IC50 and E_max_. (***ii*** and ***iii***) Only the IC50 of clone #2 is statistically higher than clone #4, while the E_max_ of clone #1, #2, and #3 are significantly higher than clone #4, suggesting that clone #4 is more sensitive to the drug treatment. (N = 3 experiments, n = 2–3 wells, Welch’s t-test, *p* < 0.05 vs. clone #4).

To investigate whether the MIA PaCa-2 clones have distinct migration rates, we utilized the scratch wound healing assay ^26,27^. Since cell proliferation contributes to wound closure in the scratch wound healing assay, to only study cell migration in this assay, we blocked cell proliferation during the process of the assay by administration of DNA synthesis inhibitor thymidine ^42,43^ in the cell culture medium 24 hours before introducing the scratch. Results indicated that wound closure was significantly faster in clone #1 and clone #2 after 48-hour culture relative to clone #3 and clone #4 (**Figure 2C,D**), suggesting the differential migration potential between the clones.

Gemcitabine is one of the first-line chemotherapeutic drugs for PDAC; it is an analog of deoxycytidine, which prevents cell growth through the inhibition of DNA synthesis ^44,45^. To assess the cytotoxic effect of gemcitabine on the MIA PaCa-2 clones, we treated the four clones with a series of gemcitabine concentrations for 3 days, followed by a luminescence cell viability assay (**Figure 2E**). Curve fitting ^28^ was performed on the luminescence data to derive the gemcitabine dose-response curves and their metrics, including the half maximal inhibitory concentration (IC50) and the least percentage of surviving cells (E_max_), as the indication of the drug potency and efficacy, respectively. In general, the IC50 of clones #1, #2, and #3 are higher than clone #4, however, only the IC50 of clone #2 is statistically different to clone #4. Meanwhile, the E_max_ of clones #1, #2, and #3 are significantly higher than clone #4. Taken together, the lower IC50 and E_max_ values for clone #4 suggest a greater sensitivity to gemcitabine compared to the other clones.

### Genomic heterogeneity between the MIA PaCa-2 clones and their stability with culture

To investigate the level of genomic heterogeneity within the MIA PaCa-2 cells, we analyzed the chromosome copy number profiles of the clones using SNPa. We have previously shown that SNPa produced similar copy number profiles as the comparative genome hybridization array ^46^, which is the gold standard for chromosome copy number profile quantification ^47^. The genomic DNA of the MIA PaCa-2 parental bulk and the single-cell clones were subjected to SNPa and the data was used to derive the copy number profiles at 1 Mb resolution (**Figure 3A**). Chromosome copy number alterations were observed across the genome of the MIA PaCa-2 clones, with distinct differences between the clones. To better visualize the copy number differences between the clones, we then derived the copy number variations (CNVs) between the MIA PaCa-2 clones and the parental bulk population (**Figure 3B**). CNVs that are unique to each clone were observed across the genome, suggesting genomic heterogeneity within the MIA PaCa-2 population.

**Figure 3.**
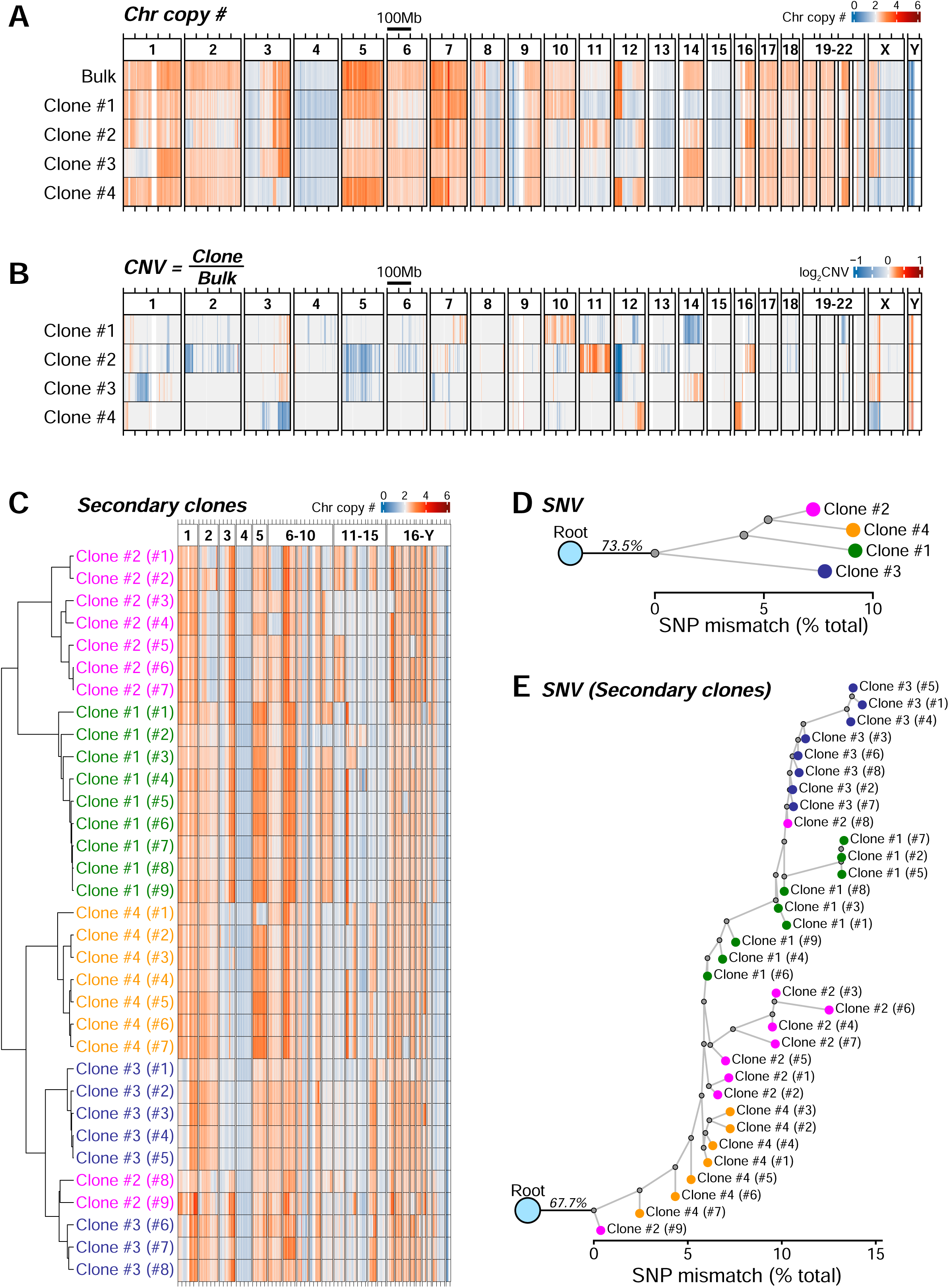
Genomic heterogeneity between the MIA PaCa-2 clones and their stability with culture. (**A**) Genomic DNA of the MIA PaCa-2 parental bulk and the single-cell clones were subjected to SNPa and the resulting data were used to derive chromosome copy numbers of the clones within 1 Mb binning windows. The heatmaps illustrate chromosome copy numbers across the whole genome, where blue indicates chromosome loss (<2) and red indicates chromosome gain (>2). (**B**) The copy number variations (CNVs) between the clones and the parental bulk were derived by dividing the copy number of each clone by the parental bulk copy number. CNVs that are unique to a clone were observed across the genome, suggesting genomic heterogeneity between the clones. (**C**) Secondary single-cell clones were derived from the primary clones #1, #2, #3, and #4, and were subjected to SNPa to derive their chromosome copy number profiles. In general, most of the secondary clones clustered together based on their respective primary clones, suggesting the genomic stability of these clones with culture. However, some differences in copy number are also observed between the secondary clones derived from the same primary clone, indicating some level of plasticity in the genome over time, even though it is less than the differences observed between the primary clones. (**D**) The SNP sequence data was used to derive the single nucleotide variations (SNVs) between the primary clones, followed by the construction of a phylogenetic tree by using the Neighbor-Joining algorithm. The tree visualizes the genomic heterogeneity between the primary clones at the single nucleotide level, covering the distance of around 10% of the total detectable SNPs (∼70,000 out of the 759,993 SNPs). (**E**) A phylogenetic tree was constructed from the SNVs between the secondary clones. Similar to the CNVs, most of the secondary clones cluster together based on their respective primary clones, but, some degree of SNVs between the secondary clones derived from the same primary clone are also observed.

To investigate the genomic plasticity of the MIA PaCa-2 cell line, secondary single-cell clones were generated from the primary clones #1, #2, #3, and #4. Subsequently, these secondary clones were subjected to SNPa analysis to determine their chromosome copy number profiles (**Figure 3C**). Hierarchical clustering analysis revealed that, overall, most secondary clones clustered together based on their corresponding primary clones. This observation suggests that the copy number profiles of secondary clones originating from a specific primary clone exhibit greater similarity to each other compared to secondary clones derived from other primary clones. This implies a certain level of genomic stability in MIA PaCa-2 cells during extended culture. However, upon closer examination of copy number profiles within secondary clones derived from the same primary clone, some variations in chromosome copy numbers were evident. This indicates a degree of genomic plasticity over time, although it is less pronounced than the differences observed between primary clones. Like the CNVs, the cellular morphology of the secondary clones resembles that of their corresponding primary clones (**Figure S2**). For instance, the spindle morphology is evident in the secondary clones derived from clone #1, while the clone #4 secondary clones exhibit a rounded morphology. Nevertheless, the appearance of elongated cells was observed in some of the clone #4 secondary clones, indicating a degree of plasticity in cellular morphology as well.

Next, to investigate genomic variations at the single nucleotide level, the SNP sequence data was utilized to identify single nucleotide variations (SNVs) among the primary clones. This information was then used to construct a phylogenetic tree using the Neighbor-Joining algorithm (**Figure 3D**). The tree visually represents the extent of SNVs among the primary clones, encompassing approximately 10% of the total detectable SNPs, equivalent to around 70,000 SNVs out of the total detectable 759,993 SNPs. Similarly, a phylogenetic tree was generated from the SNVs identified among the secondary clones (**Figure 3E**). Consistent with the CNVs, most secondary clones clustered together based on their respective primary clones. However, there were some observable SNVs among secondary clones derived from the same primary clone, suggesting a level of genomic plasticity over time.

### Transcriptomic heterogeneity between the MIA PaCa-2 clones

To investigate mRNA-level heterogeneity, RNA-seq analysis was conducted on primary clones #1, #2, #3, and #4. Uniquely regulated genes for each clone were identified through multiple pair-wise differential expression analyses against the other three clones (e.g. clone #1 compared to clones #2, #3, and #4). A gene was considered uniquely regulated when it exhibited consistent regulation (either upregulation or downregulation) in all three pair-wise comparisons (**Figure 4A-B**). The observed differential regulation of gene expression among the clones indicates transcriptomic heterogeneity within the MIA PaCa-2 population. Furthermore, we observed a general positive correlation between the direction of gene expression regulation in clone #4 and the associated CNVs (**Figure S3**), implying a potential gene dosage effect of the CNVs. To note, instances of gene expression regulation lacking corresponding CNVs were also observed, suggesting the involvement of additional regulatory mechanisms, such as epigenetic regulation.

**Figure 4.**
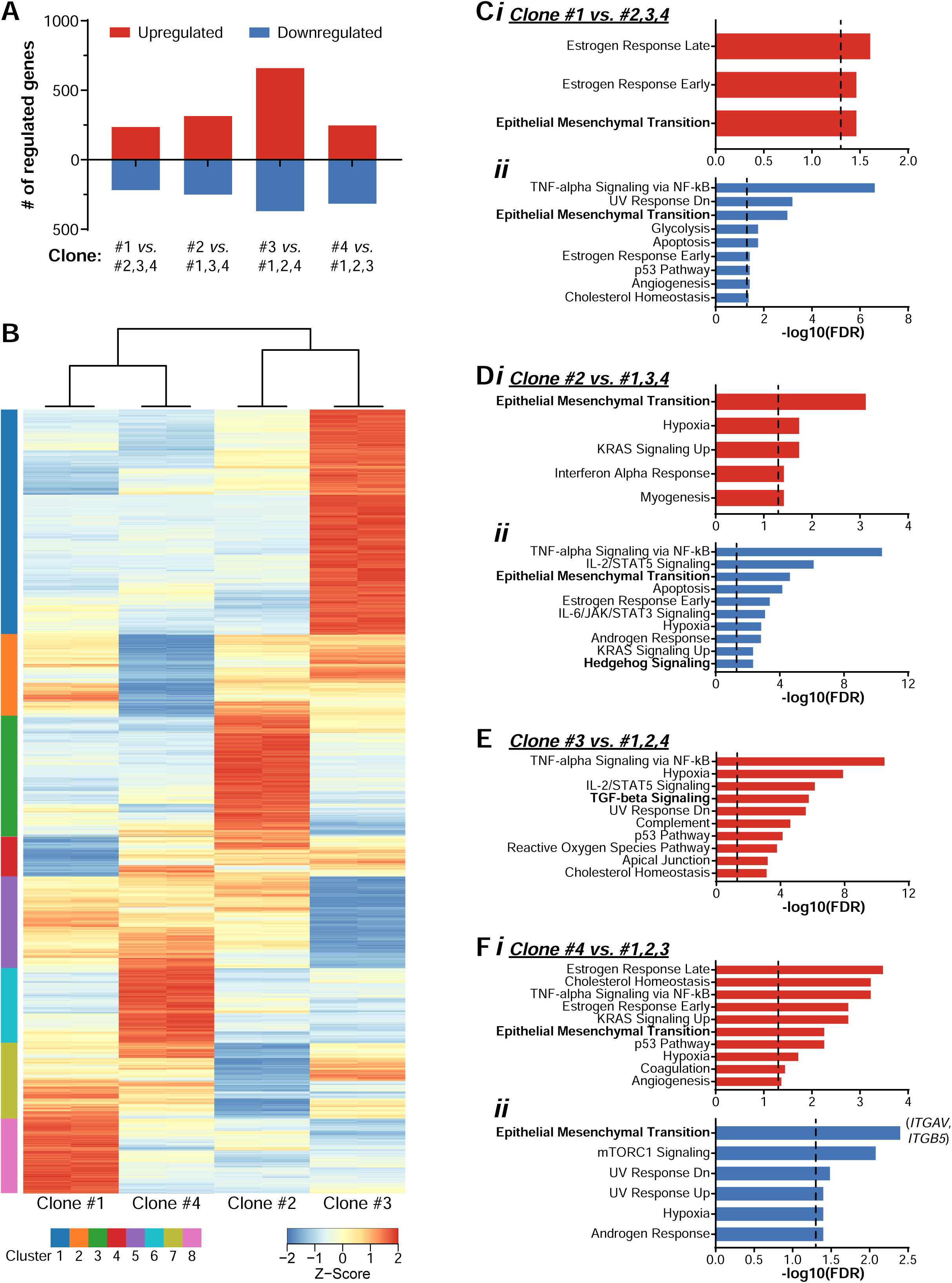
Differential gene expressions between the MIA PaCa-2 clones. The primary clones #1, #2, #3, and #4 were subjected to RNA-seq, and to identify the uniquely regulated genes, differential expression analyses were performed between a given clone against the other three clones, such as clone #1 vs. clones #2, #3, and #4. A gene is considered to be uniquely regulated when it is regulated in the same direction (upregulated or downregulated) within all of the three comparisons. (**A**) The bar plot shows the number of genes that are uniquely regulated in each of the clones. (**B**) Heatmap showing normalized read counts of the uniquely regulated genes. (**C-F**) Bar plots showing the “hallmark” gene sets that are enriched by the uniquely regulated genes of each clone. The regulated genes were subjected to gene set enrichment analysis using Enrichr and the top 10 statistically significant gene sets were selected (FDR < 0.05). The red and blue bars indicate upregulated and downregulated gene sets respectively. The dashed line on the bar plots indicates the FDR cut-off.

To examine the regulated biological processes, gene set enrichment analyses were performed on the regulated genes using Enrichr ^35^, with reference to the “hallmark” gene sets of the human molecular signatures database (MSigDB) ^36^. Various gene sets were found to be enriched across the clones (**Figure 4C-F**, **Table S1**), encompassing pathways such as estrogen response, TNF-alpha signaling, p53 pathway, hypoxia, KRAS signaling, TGF-beta signaling, JAK/STAT signaling, mTORC1 signaling, and others. This confirms the distinct transcriptomic profiles and regulations between the clones. Notably, the epithelial-mesenchymal transition gene set was enriched in nearly all clones, except for clone #3. This enrichment aligns with the observed differences in cellular morphology, which is a marker for the epithelial-mesenchymal transition process.

### *ITGAV* dictates the morphology of MIA PaCa-2 clones

In typical cell culture, the extracellular matrix (ECM) proteins within the serum are adsorbed by the culture surface and cellular adhesion is initiated by the recognition of these proteins by integrins, followed by focal adhesion complex formation and maturation. Integrins are heterodimeric proteins found across the plasma membrane that connect the cells with the ECM by binding to specific recognition sites within ECM ligands ^48,49^. Considering the importance of integrins in dictating cellular spreading and morphology, we were prompted to investigate whether any of the integrin genes are uniquely regulated between the clones, which may explain the morphological heterogeneity that we observed in Figure 1. Interestingly, *ITGAV* and *ITGB5* are listed in the epithelial-mesenchymal transition gene set that was enriched by the genes that are uniquely downregulated in clone #4 (**Figure 4Fii**, **Table S1**), the clone with the rounded morphology. Indeed, the mRNA level of *ITGAV* and *ITGB5* was the lowest for clone #4 (**Figure 5A, S4A**), and copy number loss was observed in the chromosome locations of both genes (**Figure S3**), suggesting the possibility that the expression regulation is driven by the loss of gene copy number. The lower expression level was also confirmed at the protein level through immunoblotting against ITGAV (**Figure 5B**). *ITGAV* encodes for the protein integrin αV and *ITGB5* encodes for integrin β5, which can bind to the RGD motif of fibronectin and vitronectin as heterodimers αVβ1, αVβ3, αVβ5, αVβ6, and αVβ8 ^48,49^. The depletion of *ITGAV* or *ITGB5* may be the cause of the rounded morphology of clone #4 cells.

**Figure 5.**
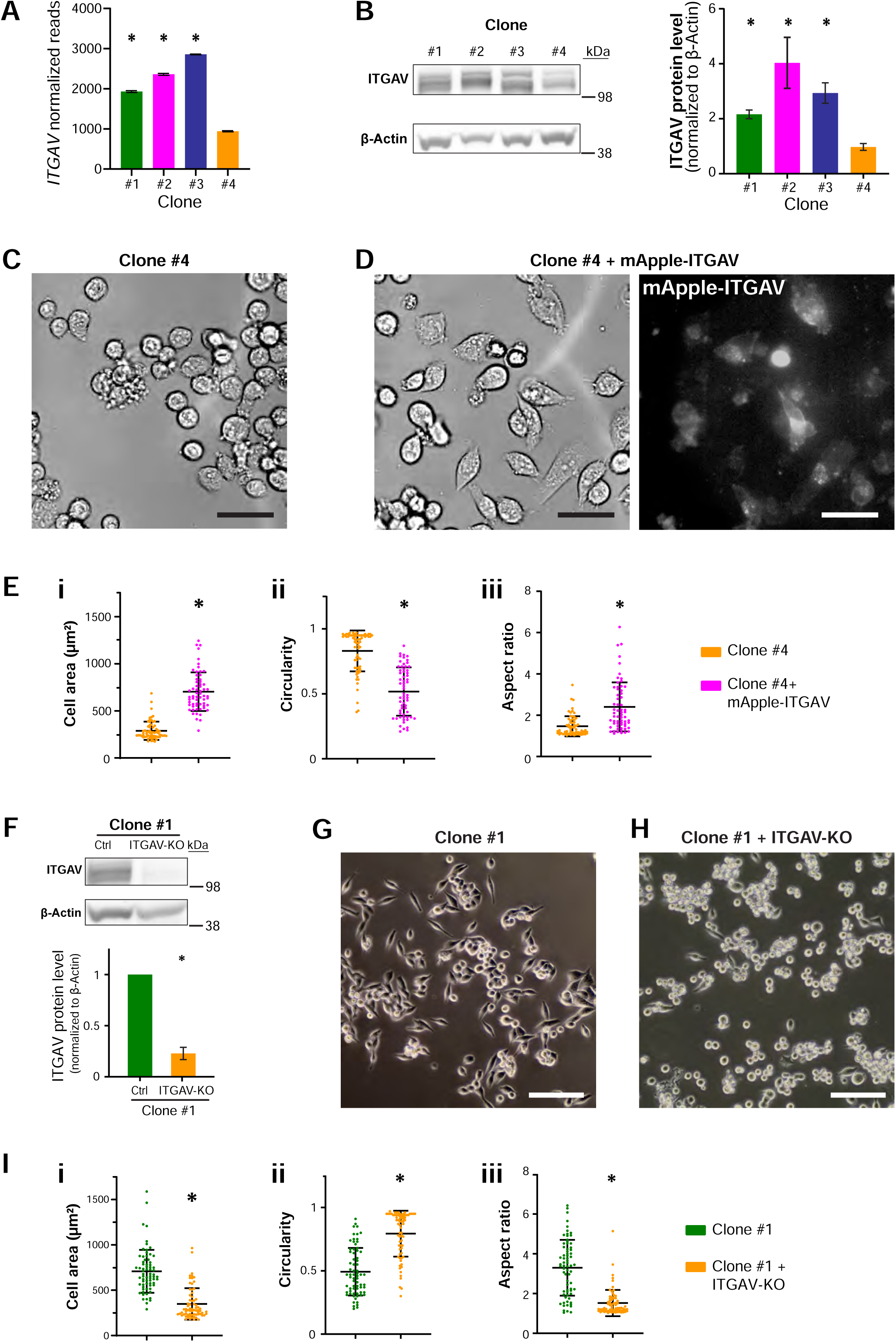
*ITGAV* dictates the cellular morphology of MIA PaCa-2 clones. (**A**) *ITGAV* read counts from the RNA-seq of the MIA PaCa-2 clones (normalized to counts per million reads, CPM), showing clone #4 had significantly decreased expression of *ITGAV* (Welch’s t-test, *p* < 0.05 vs. clone #4). (**B**) Western blot of the ITGAV protein in the MIA PaCa-2 clones (upper band) and the β-Actin protein as the loading control (lower band) (left). Quantification of the ITGAV protein level, normalized to β-Actin, showing clone #4 had a significantly lower level of the ITGAV protein (Welch’s t-test, *p* < 0.05 vs. clone #4, right). (**C** and **D**) Representative images of the MIA PaCa-2 clone #4 and clone #4 transiently transfected with mApple-ITGAV, showing that clone #4 cells were elongated when transfected with ITGAV (mApple positive) (scale bar = 50 μm). (**E**) Three cell morphometric parameters (cell area, circularity, and aspect ratio) were analyzed by ImageJ, showing significant differences between clone #4 and clone #4 expressing mApple-ITGAV (Welch’s t-test, *p* < 0.05 vs. clone #4). (**F**) Western blot showing clone #1 upon knockout of *ITGAV* significantly reduced the ITGAV protein level (normalized to β-Actin, Welch’s t-test, *p* < 0.05 vs. Ctrl). (**G** and **H**) Representative images of the MIA PaCa-2 clone #1 and clone #1 transduced with a CRISPR vector targeting *ITGAV*, showing that the majority of clone #1 cells upon depletion of *ITGAV* changed from elongated to rounded morphology (scale bar = 100 μm). (**I**) Three cell morphometric parameters (cell area, circularity, and aspect ratio) were analyzed by ImageJ, showing significant differences between clone #1 and clone #1 with the depletion of *ITGAV* (Welch’s t-test, *p* < 0.05 vs. clone #1).

To check whether ITGAV or ITGB5 is sufficient to induce morphological changes in the MIA PaCa-2 clones, we first transiently expressed mApple-ITGAV and EGFP-ITGB5 in the rounded clone #4. Unlike the EGFP-ITGB5 (**Figure S4B**), the overexpression of mApple-ITGAV induced morphological changes in the clone #4 cells, from rounded to a more elongated morphology (**Figure 5C–D**). Further quantification of the cellular morphology confirmed the significant differences between clone #4 cells with and without the expression of mApple-ITGAV (**Figure 5E**), both the cell area and the circularity of the mApple-ITGAV positive cells changed to a level similar to clone #1, while the aspect ratio increased to a degree comparable to that of clone #2 (**Figure 1C**). These results suggest that ITGAV can drive the spreading of clone #4 cells. Next, we used the CRISPR/Cas9 system to deplete the *ITGAV* expression in clone #1 (**Figure 5F**). We chose clone #1 cells because of their elongated morphology. Indeed, the knockout of *ITGAV* caused the rounding of clone #1 cells (**Figure 5G, H**). Quantification of the cell morphology parameters confirmed the significant morphological changes of clone #1 upon loss of *ITGAV* (**Figure 5I**). Taken together, these results suggested that *ITGAV* expression level can dictate the cellular morphology of MIA PaCa-2 cells.

## Discussion

Cancer undergoes cycles of clonal expansion, genetic diversification, and selection of adaptive clones, a phenomenon reminiscent of Darwin’s natural selection ^1^. Within a tumor, specific mutations provide a growth advantage to certain cancer cells, leading to the development of distinct cellular clones ^2^ and augmenting the heterogeneity within the population. This evolutionary process also provides the opportunity for the emergence of cancer cells with advantageous phenotypes, such as increased migration potential for metastasis and resistance to therapy ^3^. Multiple studies have reported the heterogeneity in PDAC tumors ^4–7^, which is also observed in PDAC models, such as patient-derived xenografts ^50^, organoids ^15,51^, and cell lines ^52–55^. Notably, genomic ^52^ and phenotype ^54,55^ heterogeneity of MIA PaCa-2 cells have been reported in separate studies. In our study, we have characterized the phenotype, genomic, and transcriptomic heterogeneity of the same clones, providing a more thorough and controlled analysis of MIA PaCa-2 population heterogeneity. Furthermore, our investigation of secondary clones also sheds light on the stability and plasticity of the MIA PaCa-2 genome and the regulation of cellular morphology.

The findings in this study showed that MIA PaCa-2 is comprised of cells with distinctive phenotypes, heterogeneous genomes, and differential transcriptomic profiles, suggesting its suitability as a model to study the underlying mechanisms behind PDAC tumor phenotypic and genomic heterogeneity. We can start such study with a single-cell clonal population, followed by subsequent cloning and genomic analysis to obtain the intrinsic genomic instability level of the cells. We can then investigate the role of the known PDAC driver genes on genomic variations by genetically modulating the corresponding genes and monitoring for any deviation from the intrinsic genomic instability level.

On a side note, it is crucial to recognize certain limitations when interpreting the findings of this study. Firstly, the resolution of our assessment of population heterogeneity is constrained by the sampling number. For the initial cellular morphology screen (Figure S1), we established 21 single-cell clones. Probability wise, our detection resolution is limited to clones that constitute more than ∼5% of the population, equivalent to around 1/21 of the population. Consequently, there is a possibility that we overlooked rare clones constituting less than 5% of the population, and this resolution could be enhanced by using higher sampling techniques, such as single-cell sequencing. Secondly, we did not characterize the epigenomic heterogeneity of the MIA PaCa-2 cells. Epigenomic heterogeneity in PDAC tumors has been reported by several studies and it was associated with tumor aggressiveness and patient prognosis ^56,57^, suggesting the critical role of the epigenetic landscape in PDAC progression. Further characterization of the MIA PaCa-2 epigenome is needed to check whether MIA PaCa-2 is a suitable model for studying the mechanisms that regulate PDAC epigenomic heterogeneity and its downstream impact on the heterogeneous transcriptomic profiles. Cancer cell lines are versatile models for studying cancer biology, enabling researchers to perform in-depth mechanistic studies at a fraction of the cost, time, and labor required by animal or human studies. However, the use of cell lines comes with various limitations, one of them being the phenotypic and genomic shift associated with decades of culture on plastic surfaces, compromising the cell line’s relevance to the original tumor ^58^. One alternative to this issue is the use of PDAC organoids derived from patient tumors ^59–65^. These organoids provide researchers with the opportunity to perform various *in vitro* mechanistic studies on a model that closely recapitulates the *in vivo* tumor phenotype, genotype, and even drug response. Furthermore, several studies have shown genomic and transcriptomic heterogeneity within PDAC organoids ^15,51^, suggesting that organoids are suitable for studying genomic instability in PDAC.

## Data availability

The genomic data will be made available in Sequence Read Archive and Gene Expression Omnibus.

## Acknowledgment

The authors would like to thank Drs. Cynthia Vied and Yanming Yang at Florida State University’s Translational Science Laboratory for running the RNA-seq libraries and Dr. Beth Alexander at Florida State University’s Flow Cytometry Laboratory for helping isolate the MIA PaCa-2 clones. We are grateful for the great service from the team at Florida State University’s Biology Core Facilities. The authors would like to thank Dr. Terra Bradley for the careful editing of the manuscript. The authors thank all the lab members in Dr. Jerome Irianto’s and Dr. Yue Julia Wang’s laboratories for their valuable suggestions and assistance. The authors in this study were supported by startup funds from Florida State University, awards from the Florida Department of Health’s Bankhead-Coley Cancer Research Program (award number 21B11) and Live Like Bella Pediatric Cancer Research Initiative (award number 23L06), and a collaborative Trans-Network Project linked to the Physical Science-Oncology Project 5U01 CA214282 from the National Cancer Institute of the National Institutes of Health.

---

**Figure S1.**
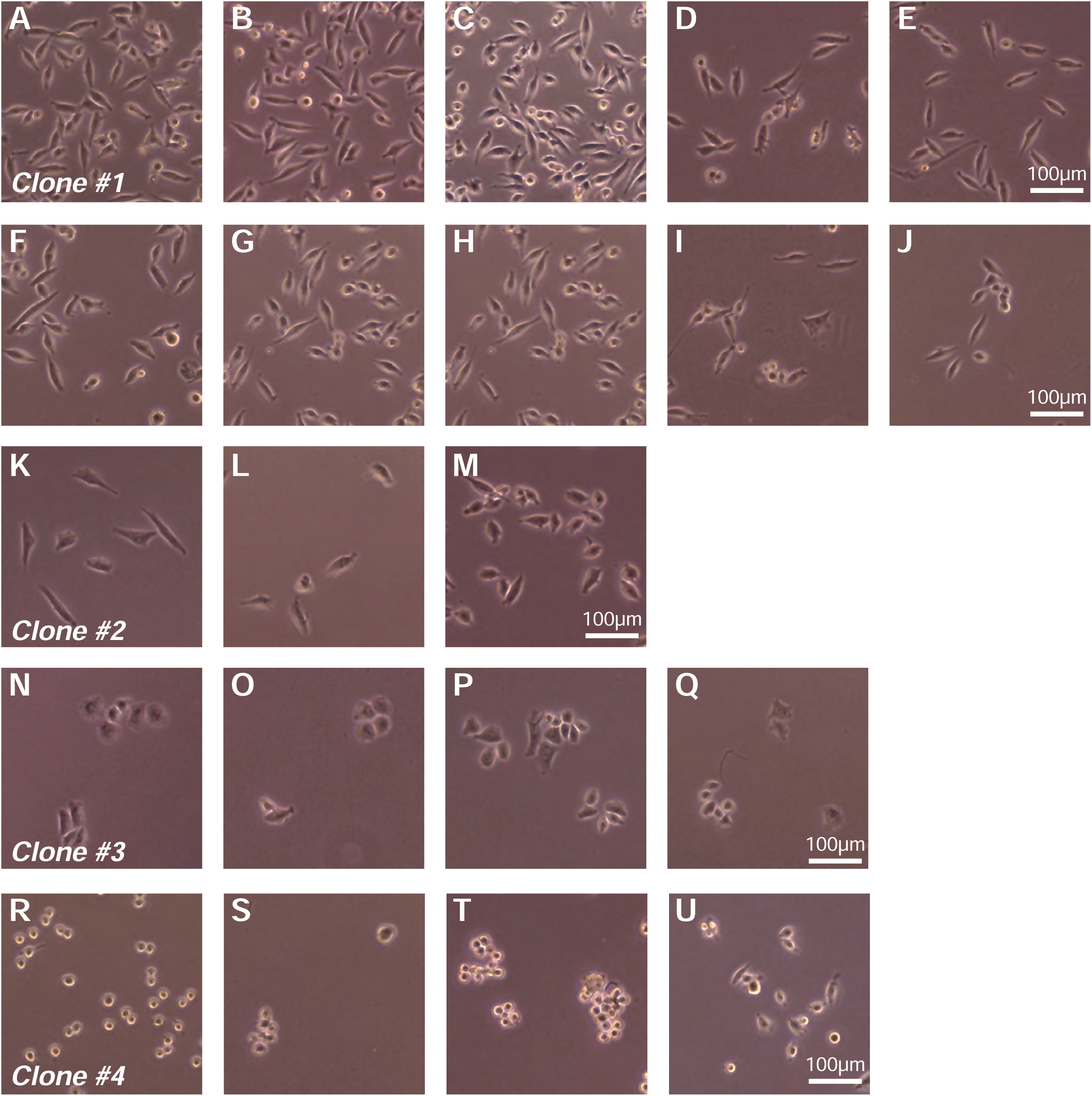
Representative images of the 21 single-cell clones established from the MIA PaCa-2 bulk population, showing heterogeneous cellular morphology. A = Clone #1, K = Clone #2, N = Clone #3, and R = Clone #4.

**Figure S2.**
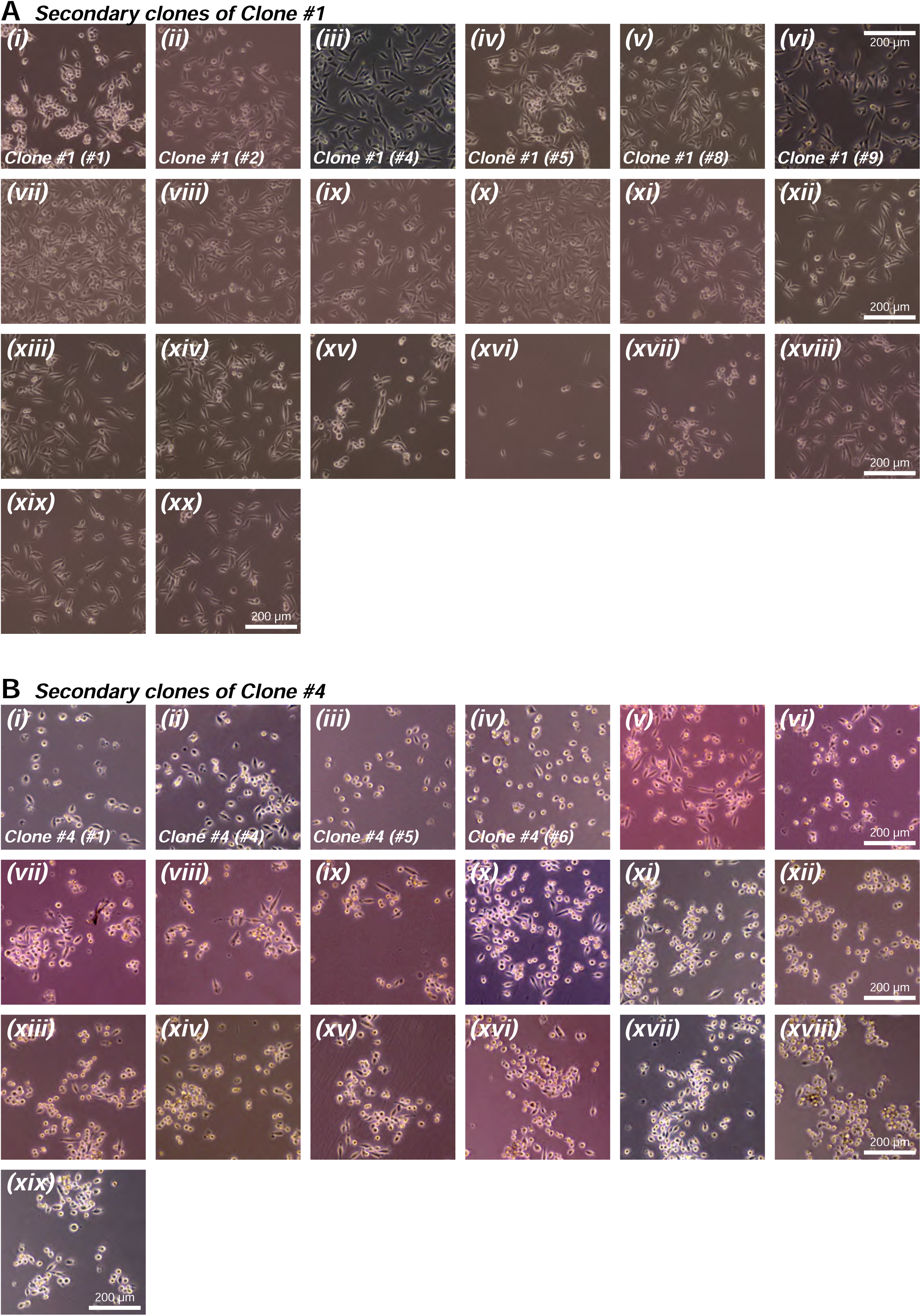
(**A**) Representative images of the 20 secondary single-cell clones derived from the primary clones #1. Most of the secondary clones have spindle morphology, resembling the parental clone #1 cellular morphology. (**B**). Representative images of the 19 secondary single-cell clones derived from the primary clone #4. Most of the secondary clones have rounded morphology, however, the emergence of more elongated cells can be observed in some clones, such as in (***ii***, ***v***, and ***x***).

**Figure S3.**
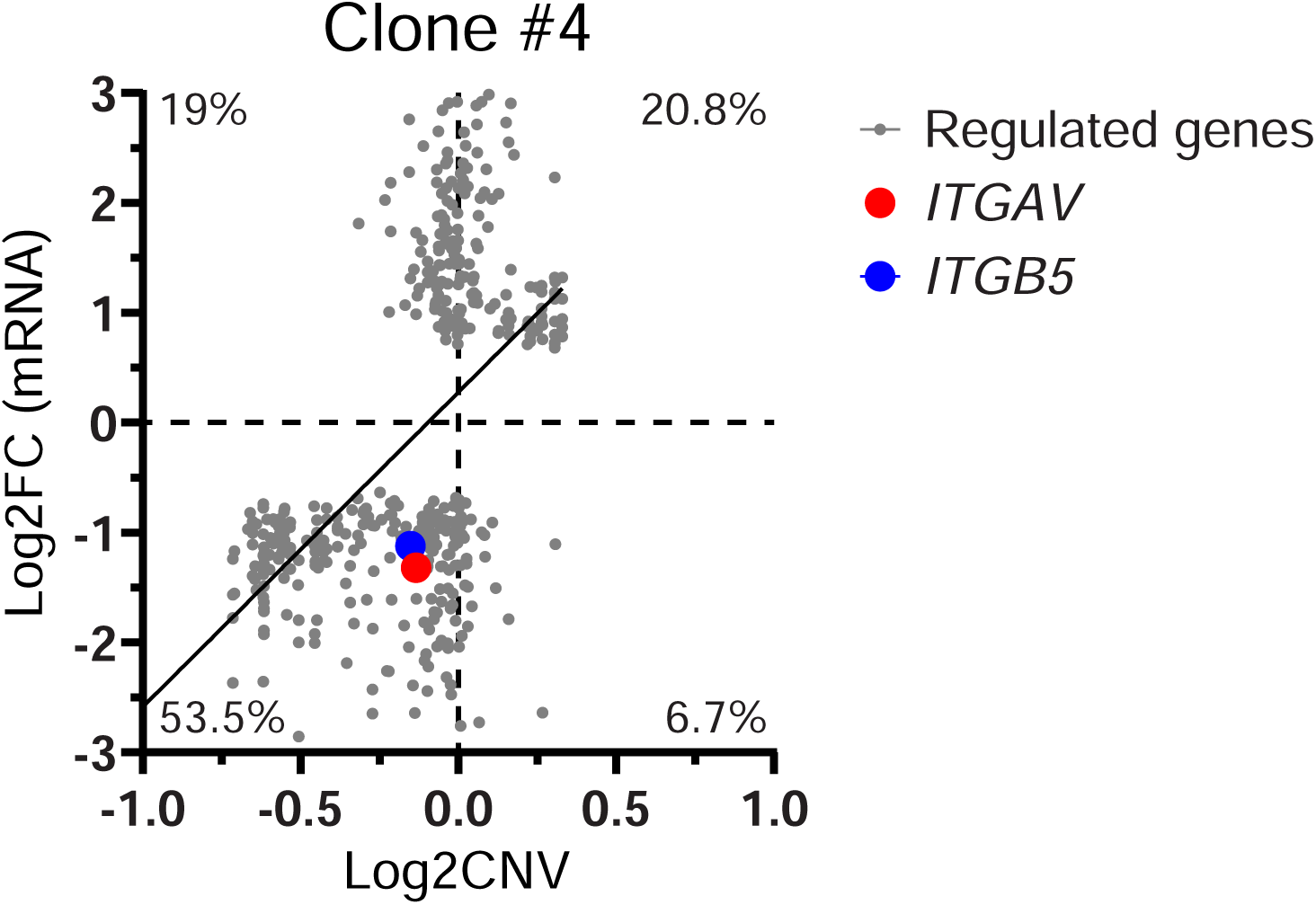
For the genes that are uniquely regulated in primary clone #4, the gene expression fold change was correlated to the copy number variation at the corresponding chromosome position. The plot shows an overall positive correlation between the two, suggesting a gene dosage effect. However, gene expression regulations that are absent of copy number variations were also observed, indicating the involvement of other regulatory mechanisms, such as epigenetic regulation.

**Figure S4.**
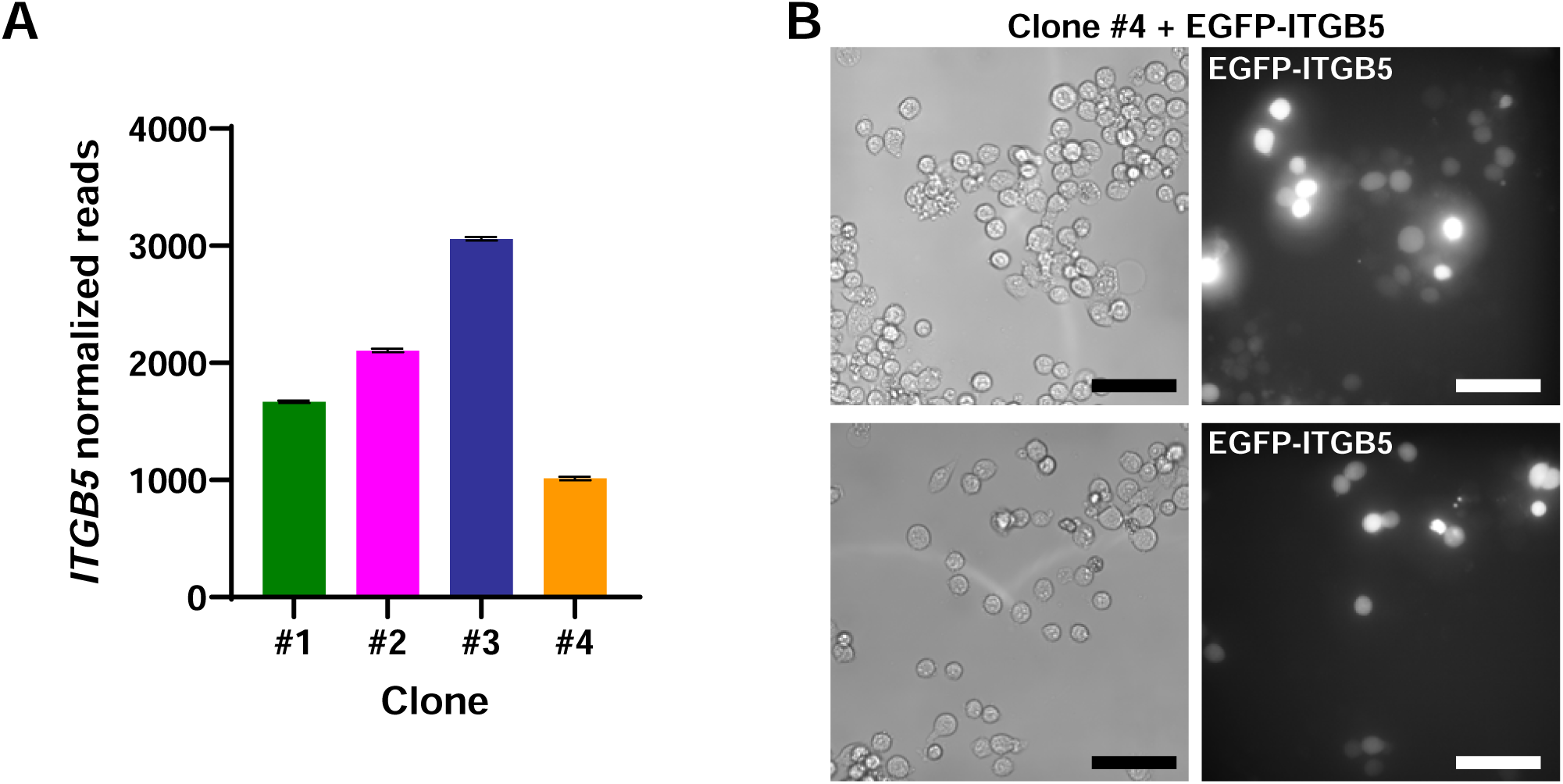
(**A**) *ITGB5* read counts from the RNA-seq of the MIA PaCa-2 clones (normalized to counts per million reads, CPM), showing clone #4 had significantly lower expression of *ITGB5* (Welch’s t-test, *p* < 0.05 vs. clone #4). (**B**) Representative images of primary clone #4 cells transiently transfected with EGFP-ITGB5, showing that ITGB5 transfected cells remained rounded, i.e. no morphological changes after ITGB5 overexpression (scale bar = 50 μm).

